# IRON MAN, a ubiquitous family of peptides that control iron transport in plants

**DOI:** 10.1101/351635

**Authors:** Louis Grillet, Ping Lan, Wenfeng Li, Girish Mokkapati, Wolfgang Schmidt

**Author notes:** These authors contributed equally to this work (Louis Grillet, Ping Lan). Correspondence and requests for materials should be addressed to W.S.

## Abstract

Iron (Fe) is an essential mineral nutrient which severely affects the growth, yield and nutritional quality of plants if not supplied in sufficient quantities. We here report that a short C-terminal amino acid sequence consensus motif (IRON MAN; IMA) conserved across numerous, highly diverse peptides in angiosperms, is essential for Fe uptake in plants. Overexpression of the IMA sequence in *Arabidopsis* induced Fe uptake genes in roots, causing accumulation of Fe and manganese in all plant parts including seeds. Silencing of all eight *IMA* genes harbored by the *Arabidopsis* genome abolished Fe uptake and caused severe chlorosis; increasing the Fe supply or overexpressing *IMA1* restored the wild-type phenotype. *IMA1* is predominantly expressed in the phloem, preferentially in leaves, and reciprocal grafting showed that IMA1 peptides in shoots positively regulate Fe uptake in roots. *IMA* homologs are highly responsive to the Fe status and functional when heterologously expressed across species. IMA constitutes a novel family of peptides which are critical for the acquisition and cellular homeostasis of Fe across land plants.

Although iron (Fe) is one of the most abundant elements on earth, the extremely low activity of free Fe in most soils often severely restricts its uptake, making Fe deficiency a common nutritional disorder in plants. In human populations, insufficient dietary Fe intake resulting from low Fe concentrations in edible plant parts is the cause of Fe deficiency-induced anemia (IDA), affecting more than one billion people worldwide, particularly in areas where Fe supply depends mainly or entirely on plants^1^. Understanding how plants regulate the uptake and distribution of Fe is thus mandatory to produce Fe-enriched germplasms and combat IDA.

Plants have evolved multifaceted strategies to acquire Fe from soils^2,3^. Rice *(Oryza sativa)* and other graminaceous species take up Fe after secretion of Fe^3+^-binding phytosiderophores (PS) and subsequent uptake of the Fe^3+^-PS complex via an oligopeptide transporter of the YSL family (Strategy II)^2,4,5^. *Arabidopsis* and all non-grass species employ a reduction-based Fe acquisition mechanism (Strategy I), in which Fe^3+^ is first reduced by the Fe^3+^-chelate reductase FRO2. The reduced Fe^2+^ is then transported across the plasma membrane by the ZIP family transporter IRT1^6–8^. Solubilization of recalcitrant Fe pools is facilitated by P-type ATPase-driven proton extrusion^9^. The two Fe acquisition strategies are thought to be mutually exclusive^2^. However, rice possesses an Fe^2+^ uptake system^10^ and, similar to the PS-system of grasses, *Arabidopsis* and other non-graminaceous species secrete Fe^3+^-mobilizing coumarins^11–15^, indicating that the two mechanisms share analogous components.

In *Arabidopsis*, the bHLH-type transcription factors PYE and FIT control non-overlapping subsets of genes involved in the acquisition and cellular homeostasis of Fe^16,17^. FIT forms heterodimers with the 1b subgroup bHLH proteins bHLH38, bHLH39, bHLH100 and bHLH101^18,19^ Both FIT and PYE are directly activated by bHLH34, bHLH115 and bHLH105 (ILR3)^20,21^. The abundance of bHLH104 and bHLH105 is regulated by the Fe-binding E3 ligase BTS^22^. In rice (*Oryza sativa*), OsIRO2, an ortholog of AtbHLH38/39, regulates the Fe^3+^-PS transporter OsYSL15^23^ but not the uptake of Fe^2+^ via OsIRT1^24^. Two orthologs of BTS, OsHRZ1 and OsHRZ2, negatively regulate Fe uptake presumably via OsIRO2, OsIRO3, an ortholog of PYE^25,26^, and OsPRI1, an ortholog of bHLH105^27^.

The regulation of root Fe acquisition by shoot-derived signals has been demonstrated more than two decades ago using a graft-transmissible, Fe-accumulating trait of the pea mutant *dgl*^28^. In *Arabidopsis*, evidence for such signals reside in the enhanced Fe uptake of *frd3*^29^ and *opt3*^30,31^ mutants, respectively defective in root-to-shoot transport and phloem loading of Fe. The nature of the long-distance signal that conveys information of the Fe status of leaves to the roots is a long-standing enigma in Fe research. In nitrogen-deprived roots, members of a family of 15 amino acid C-terminally Encoded Peptides (CEPs) activate the leucine-rich repeat receptor kinase CEPR^32^. CEPR phosphorylates the phloem-localized class III glutaredoxin CEPD1^33^, which subsequently acquires the ability to exit the phloem and migrate to the endodermis. How CEPD1 triggers the expression of the nitrate uptake transporters NRT1.1, NRT2.1 and NRT3.1 remains elusive.

Here, we describe the discovery of a novel peptide family expressed in the phloem that presumably act as a phloem-mobile signal to control Fe uptake in *Arabidopsis* and, possibly, constitutes a common component of Fe signaling across Magnoliophyta. Members of this family harbor a 17-amino acid C-terminal consensus motif highly conserved across angiosperms that is necessary and sufficient for Fe uptake from the soil.

## Results

Similarities in the proteins controlling cellular Fe homeostasis between rice and *Arabidopsis* suggest signaling nodes that are conserved across species. To discover novel components with critical function in Fe homeostasis, we searched for common sequence motifs in Fe-responsive proteins of unknown function in rice and *Arabidopsis*, two species with well-explored Fe deficiency responses. To this end, we mined expression data of Fe-responsive genes that showed greater than 5-fold changes in transcript abundance in response to Fe deficiency. Sequences of 14 rice and *Arabidopsis* genes encoding proteins of unknown function^34,35^ were screened for common sequence motifs (Supplementary Table 1). The C-terminal amino acid sequence G-D-D-D-D-x(1,3)-D-x-A-P-A-A was found to be conserved in two *Arabidopsis* (At1g47400 and At2g30766) and two rice proteins, corresponding to LOC_Os01g45914 and to a non-annotated transcript encoded by a gene located between LOC_Os07g04910 and LOC_Os07g04930 that we designated LOC_Os07g04920 (probe sets Os.12430.1.S1_at and Os.48053.1.A1_at).

Transgenic plants over-expressing At1g47400 under the control of the CaMV 35S promoter displayed necrotic spots on the leaves, resembling Fe toxicity symptoms (Fig. 1a). Using Perls’ and Perls’-DAB Fe staining, we observed high Fe levels in leaves, in the stele, and in embryos (Fig. 1a). In histological sections of rosette leaves from the wild type, Fe was detected in xylem vessels, nuclei, and as a diffuse, homogenous signal in plastids (Fig. 1b-e). By contrast, in rosettes of 35Spro::At1g47400_cDNA_ (IMA1c Ox) lines, xylem and nuclei were more heavily stained and plastids were scattered with numerous Fe-rich granules resembling ferritins (Fig. 1f-i)^36^. Fe accumulation in the apoplast around shrunk cells was evident at necrotic regions (Fig. 1j, k), confirming that necrosis was associated with excess Fe accumulation. Mineral nutrient analysis of IMA1c Ox plants by ICP-MS revealed dramatically increased levels of Fe, zinc (Zn) and manganese (Mn). In rosette leaves, an up to 10-fold increase in Fe, a 6-fold increase in Mn, a 4-fold increase in Zn but no change in copper (Cu) concentration was observed when compared to the wild type (Fig. 1l; Supplementary Fig. 1b). Importantly, the seed Fe concentration was increased two-to three-fold in transgenic lines. Seed yield was largely unaffected and only slightly reduced in two overexpression (Ox) lines (#0-8 and #2-1; Supplementary Fig. 1c).

**Fig. 1.**
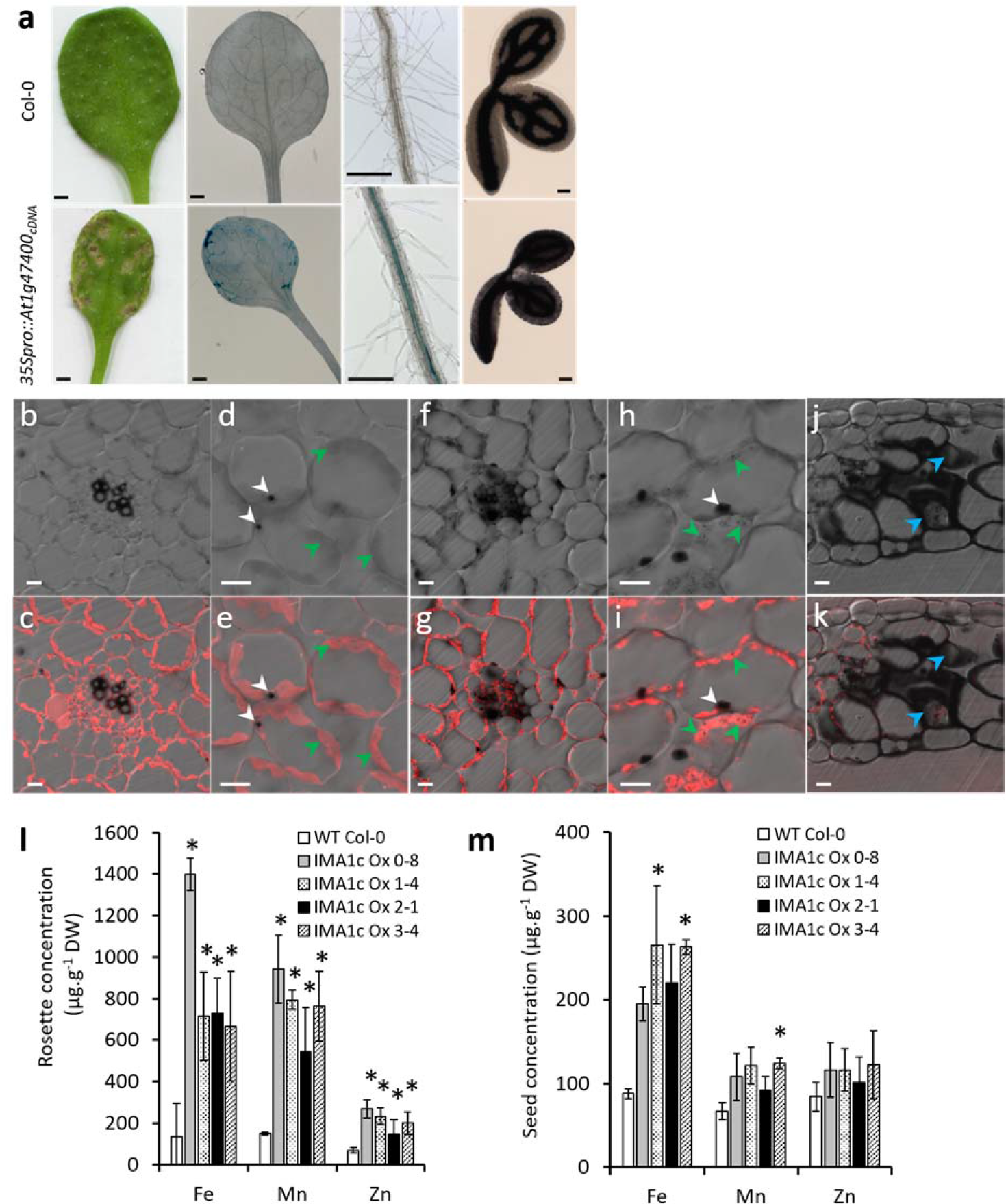
Overexpression of *IMA1* triggers Fe, Mn and Zn accumulation in *Arabidopsis*. (a) Leaves, roots and embryos of Col-0 (upper panel) and 35Spro::IMA1_cDNA_ (IMA1c Ox) plants (lower panel). Leaves of IMA1 Ox lines show necrotic spots (‘bronzing’) due to Fe overaccumulation. Leaves and roots were stained with Perls’ reagent; embryos were stained with Perls’ reagent plus DAB to reveal Fe accumulation. (b-k) Fe localization in sections of resin-embedded leaves stained with Perls’-DAB. (b-e) Leaves of wild-type plants. (f-k) Leaves of IMA1 Ox plants. (b-i) Fe accumulation in vascular tissues (b, c, f, g) and subcellular Fe localization in mesophyll cells (d, e, h, i). High Fe concentrations were observed in nuclei, nucleoli and plastids. (h, i) Dot-shaped structures are visible only in plastids of IMA1 Ox lines. (j, k) Necrotic spots in IMA1 Ox leaves. Upper panel, differential interference contrast (DIC) pictures; lower panel, DIC and autofluorescence overlap. White arrows denote nuclei, green arrows indicate plastids, blue arrows point to shrunk necrotic cells. (l) Quantification of Fe, Mn and Zn in seeds and (m) rosette leaves by ICP-MS. Results are means ± SE (n = 3 sets of 3 plants). Stars indicate significant difference to control plants (Duncan test, *P* ≤ 0.05). Scale bar = 500 μm for leaves and roots, 50 μm for embryos, 10 μm for histological sections.

Owing to the observed accumulation of Fe and Mn caused by the over-expression of At1g47400, we designated genes encoding peptides that contain the G-D-D-D-D-x(1,3)-D-x-A-P-A-A consensus motif *IRON MAN (IMA)*. The *Arabidopsis* genome harbors eight *IMA* genes (Supplementary Fig. 2a), which are all responsive to the Fe supply (Supplementary Fig. 2b). *AtIMA1* (At1g47400), *AtIMA2* (At1g47395) and *AtIMA3* (At2g30766) are highly expressed in both leaves and roots of Fe-deficient plants^35,37^. By contrast, *AtIMA4*, which we designated At1g47402, *AtIMA5* (designated At1g47406), *AtIMA6* (designated At1g47407), *AtIMA7* (designated At2g44744), and *AtIMA8* (designated At1g47401) are expressed at lower levels and are not included in the TAIR10 genome annotation (Supplementary Table 2).

*IMA2* shares 82% sequence identity with *IMA1* and is organized as a tandem repeat. IMA1 and IMA3 share only sequence identity within the IMA motif. To uncover possible functional diversity among IMA peptides, we generated also transgenic lines overexpressing *IMA3*. Growth of both IMA1 Ox and IMA3 Ox lines appeared to be negatively correlated with the Fe concentration (Fig. 2a; Supplementary Fig. 3a,c). No significant growth penalty of the IMA Ox lines was observed in the absence of Fe (Fig. 2a; Supplementary Fig. 3a). Importantly, when grown on media with limited Fe availability due to immobilization of Fe by using ferric chloride as an Fe source at neutral pH (navFe, 10 μM FeCl_3_ at pH 7), rosettes of most of the IMA Ox lines had higher chlorophyll concentrations and contained significantly more Fe than control plants overexpressing EYFP (Supplementary Fig. 3b,c), indicating increased ability to acquire Fe from recalcitrant Fe pools.

**Fig. 2.**
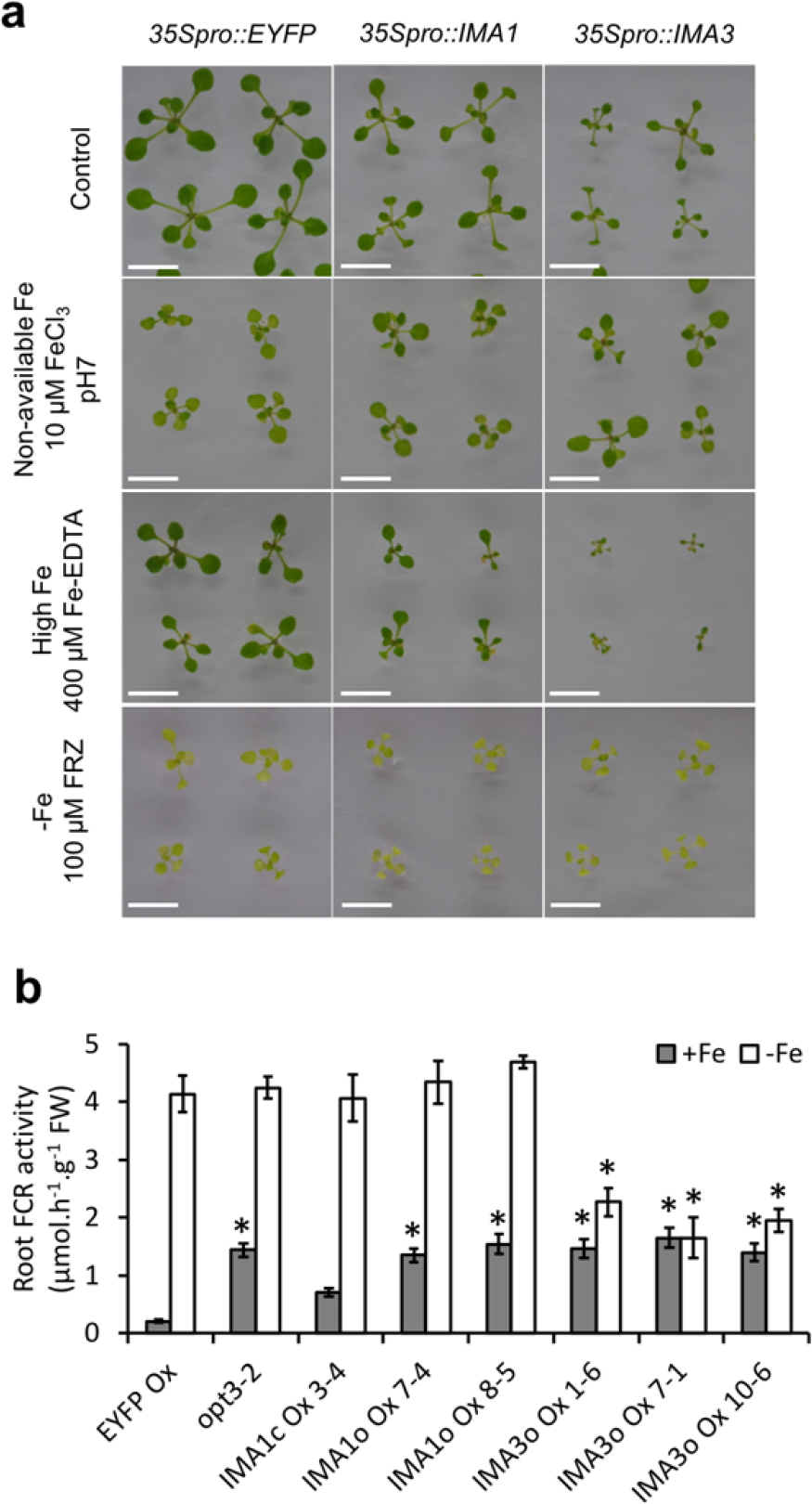
Phenotypic characterization of transgenic plants with altered expression of *IMA* genes. (a) Thirteen-day-old IMA1 Ox and IMA3 Ox plants grown under various Fe regimes. (b) Root ferric-chelate reductase (FCR) activity of plants grown for three days on Fe-replete and Fe-free media (n = 8 sets of 5 roots). Control, Estelle and Somerville (ES) media containing 40 μM FeEDTA; Non-available Fe, ES media containing 10 μM FeCl_3_, pH 7; −Fe, ES media without added Fe and supplemented with 100 μM FerroZine (FRZ); IMAlc Ox: 35Spro::AtIMA1_cDNA_; IMA1o Ox: 35Spro::AtIMA1_ORF_; IMA3o Ox: 35Spro::AtIMA3_ORF_. Results show means ± SE. Stars indicate significant difference to control plants grown under the same conditions (Duncan test, *P* ≤ 0.05). Scale bar = 1cm.

Overexpression of *IMA1* significantly increased root ferric chelate reduction (FCR) rates of plants grown under Fe-sufficient conditions; no difference in FCR activity to control plants was observed when the plants were grown on Fe-deplete media (Fig. 2b). Similar to what has been observed for IMA1 Ox plants, in Fe-sufficient IMA3 Ox lines FCR activity was constitutively increased (Fig. 2b). However, in contrast to IMA1 Ox and control plants, IMA3 Ox lines failed to further increase their FCR activity upon transfer to Fe-deplete media. Judging from the similar leaf Fe concentration of IMA1 Ox and IMA3 Ox lines, this phenotypic dissimilarity was not caused by a different Fe status of the two genotypes (Supplementary Fig. 3c). It can thus be assumed that under Fe-deficient conditions the exact role of IMA peptides in Fe homeostasis may differ among IMA family members.

Peptides harboring IMA motifs are present in the genomes of all Magnoliophyta sequenced so far including the basal angiosperm *Amboretta trichopoda*, demonstrating conservation of IMA in the flowering plant lineage. Based on the available genomic data, we identified 132 genes encoding putative IMA sequences in 29 plant species (Supplementary Table 2). This information was used to refine the IMA consensus motif (Fig. 3a). We failed to detect IMA-encoding sequences in the genomes of gymnosperms, ferns, algae or fungi, suggesting that IMA emerged at an early stage of angiosperm evolution. All IMA motif-containing genes are either unannotated or annotated as encoding unknown proteins. Notably, putative IMA homologs are among the most Fe-responsive genes in both roots and leaves of species for which data on Fe deficiency-induced changes in transcriptional profiles are available (see Supplementary Table 2 for gene IDs); *e.g.* tomato^38^ (designated *SlIMA1*), rice^34^ (designated *OsIMA1* and *OsIMA2*) and soybean^39^ (designated *GmIMA1-5*). Amino acid alignments of the encoded peptides show no sequence similarity except for the conserved IMA sequence (Fig. 3b). Alignment of the amino acid sequences of all IMA-encoding genes and a phylogenetic tree inferred from the computed sequences are shown in Supplementary Fig. 4.

**Fig. 3.**
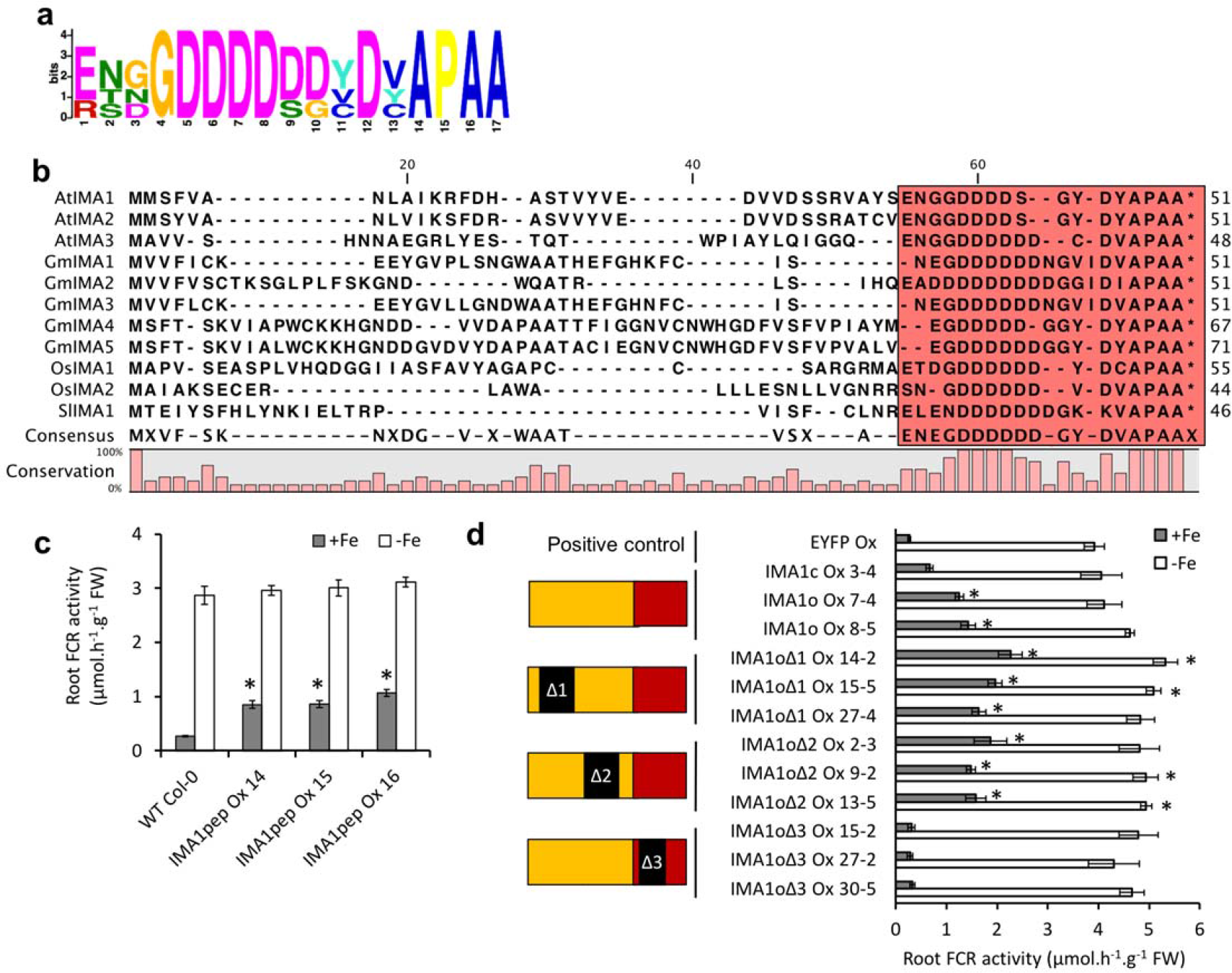
The C-terminal amino acid consensus motif is critical for IMA1 function. (a) Logo of the motif inferred from the sequences of 132 putative IMA peptides (Supplementary Table S2). (b) Amino acid alignment of the putative IMAs identified in transcriptomes of *Arabidopsis thaliana* (AtIMA), soybean (GmIMA), rice (OsIMA) and tomato (SlIMA). (c) Root FCR activity of plants expressing a peptide corresponding to the 17 C-terminal amino acids of AtIMA1 plus a N-terminal methionine residue (n = 5 sets of 5 roots). (d) Root FCR activity of plants overexpressing *IMA1* and mutated versions of the ORF harboring deletions of various parts of the protein. Results are means ± SE (n = 6 sets of 5 roots). Stars indicate significant difference to control plants grown under the same conditions (Duncan test, *P* ≤ 0.05). IMA 1c Ox, 35Spro::AtIMA 1 cdna; IMA1o Ox, 35Spro::AtIMA1ORF. IMA3o, 35Spro::AtIMA3_ORF_; IMA1pep Ox, 35Spro::IMA1pep.

**Fig. 4.**
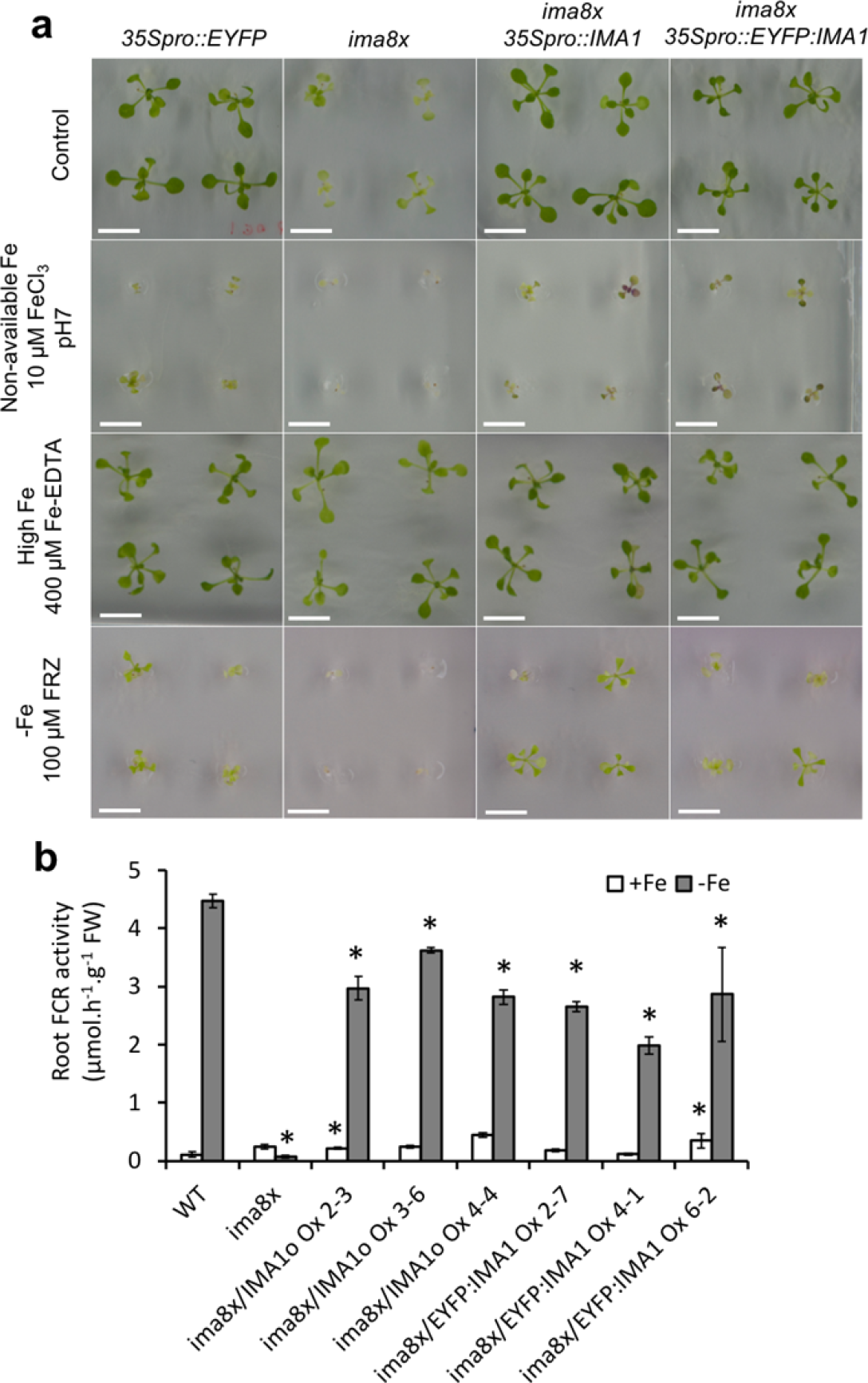
Silencing of eight *IMA* genes by CRISPR-Cas9 gene editing and complementation with *IMA1 and EYFP.IMA1.* (a) Gene editing effects on octuple *ima8x* mutants grown under various Fe regimes. Control, Estelle and Somerville (ES) media containing 40 FeEDTA; Non-available Fe, ES media containing 10 μM FeCl_3_, pH 7; -Fe, ES media without added Fe and supplemented with 100 μM FerroZine (FRZ). (b) Root ferric chelate reductase (FCR) activity (n = 6 sets of 5 roots) of wild-type plants (WT) and *ima8x* mutants grown for three days on Fe-replete and Fe-free media. Results are means ± SE. Stars indicate significant difference to control plants grown under the same conditions (Duncan test, *P* ≤ 0.05). *ima8x*, genes with silencing mutations in all eight *IMA* genes (Supplementary Fig. 7-8); ima8x/IMA1o Ox, *ima8x* plants expressing a 35Spro::IMA1 construct; ima8x/EYFP:IMA1 Ox, *ima8x* plants expressing a 35Spro::EYFP:IMA1 construct. Scale bar = 1cm.

To verify the supposition that the conserved motif is the functional part of IMA peptides, we produced transgenic plants overexpressing the 17 C-terminal amino acids of IMA1 preceded by a start codon. Under standard growth conditions, plants expressing this construct exhibited constitutively induced FCR activity similar to IMA1 Ox lines (Fig. 3c). Next, we overexpressed chimeric IMA1 ORFs harboring various deletions in the regions coding the variable N-terminus (IMA1_o_Δ1 and IMA1_o_Δ2) or the C-terminal motif (IMA1_o_Δ3). Consistent with the assumption that the C-terminal consensus sequence is critical for IMA function, plants overexpressing IMA1_o_Δ1 and IMA1_o_Δ2 developed bronzing spots and exhibited constitutively activated FCR activity, while plants expressing IMA1OA3 showed root FCR rates that were not significantly different from those of control plants (Fig. 3d). All transgenic lines expressed their respective IMA1 ORF version at comparable levels (Supplementary Fig. 5). Based on these data, we concluded that the IMA motif in IMA1 is necessary and sufficient to induce Fe uptake in roots. This result implies that there is a strong likelihood of redundancy between the eight *IMA* genes in *Arabidopsis*.

To further clarify the role of IMA1 in Fe uptake, we attempted to silence IMA1 and its close seqlog IMA2 using an artificial microRNA construct (Supplementary Fig. 6). Plants with decreased expression of both genes grew as big and healthy as control plants in all conditions tested (Supplementary Fig. 6d), and their ability to induce Fe deficiency was not affected (Supplementary Fig. 6c). To overcome putative genetic redundancy among the IMA genes, we silenced all eight *IMA* genes using CRISPR-Cas9 genome editing (Fig. 4). The generated octuple mutant (*ima8x*) carried two large deletions on chromosome 1 in the 10 kb region, containing the *IMA1-IMA6* and *IMA8* loci (Supplementary Fig. 7). The ORF of *IMA3* was subjected to a deletion in the start codon, *IMA7* contained a single nucleotide insertion resulting in a frameshift (Supplementary Fig. 8). When grown on Fe-replete media, *ima8x* plants were very small, extremely chlorotic (Fig. 4a, Supplementary Fig. 9a,b), and died within few days after transfer to soil. This phenotype was exacerbated on media with no or low available Fe and fully rescued by growing the plants on high Fe media. Induction of root FCR activity upon transfer to Fe-deplete medium was completely abolished in *ima8x* plants (Fig. 4b), suggesting that the hypersensitivity of *ima8x* plants to Fe deficiency was due to impaired Fe uptake. Consistent with the hypothesis of pronounced redundancy between *IMA* genes, overexpression of *IMA1* and *EYFP:IMA1* in the *ima8x* background restored the growth, chlorophyll content, and the FCR induction capacity almost to wild-type levels (Fig. 4a-b; Supplementary Fig. 9a-b).

To investigate the biological function of IMA1, we conducted RNA-seq transcriptome analyses of leaves and roots from Fe-deficient and Fe-sufficient IMA1 Ox plants. Genes that were differentially expressed (DEGs) between IMA1 Ox and control plants were compared with Fe deficiency DEGs of wild-type plants that were mined from previously published transcriptome data^35,11^ (Supplementary Fig. 10; Supplementary Dataset 1). Sequencing results were confirmed in other IMA1 Ox lines by qRT-PCR for a subset of genes (Supplementary Fig. 11). In roots of Fe-sufficient IMA1 Ox plants genes encoding regulators of the Fe deficiency response (e.g. the subgroup 1b bHLH proteins *bHLH38*, *bHLH39*, *bHLH100* and *bHLH101*) and proteins involved in the uptake (*FRO2* and *IRT1)* or distribution of Fe (*NAS1*, *NAS2*, and *FRD3*)^40,41^ were strongly induced in the transgenic lines (Supplementary Fig. 10a,e). Genes encoding proteins important for Fe storage, *e.g.* the ferritins FER1 and FER3^42^ and the vacuolar Fe transporters VTL1, VTL2, and VTL5^43^, were upregulated in IMA1 Ox and downregulated in control plants (Supplementary Fig. 10a,e).

In leaves, the expression profile of IMA1 Ox plants was strikingly different from that observed in roots (Supplementary Fig.10c, d, f). Genes encoding subgroup 1b bHLH proteins and other highly Fe-responsive genes were not differentially expressed between IMA1 Ox and wild-type plants. By contrast, ferritins, downregulated in Fe-deficient wild-type leaves, as well as genes involved in long-distance circulation and seed loading (*NAS3*^40^ and *YSL1*^44^), were upregulated in leaves of IMA1 Ox. It thus appears that in roots, overexpression of *IMA1* triggers a pronounced Fe-deficiency response and promotes root-to-shoot translocation of Fe, while leaves respond with the induction of genes involved in counteracting Fe-excess (Supplementary Fig.10f). This profile is reminiscent to that of the *opt3* mutant, which displays an upregulated Fe deficiency response in roots due to compromised shoot-to-root signaling^30–31^. Leaves of *opt3* plants display a transcriptional profile consistent with unimpaired local Fe sensing but compromised systemic signaling, leading to a constitutively activated Fe-deficiency response^45^.

In control plants, induction of all but one *(IMA4) IMA* genes was more pronounced in leaves than in roots (Supplementary Fig. 2). Higher transcript levels in leaves compared to roots were also observed for all putative rice and soybean *IMA* homologs^34,39^. Translatome profiling of *Arabidopsis* found *IMA1* specifically translated in the phloem^46^, prompting us to investigate whether IMAs control shoot-to-root signaling of Fe-deficiency. In transgenic lines expressing a *promIMA1::EYFP* construct, fluorescence was predominantly observed in the phloem of roots (Fig. 5a-d) and leaves (Fig. 5e,f), suggesting that IMA1 itself could constitute a mobile signal.

**Fig. 5.**
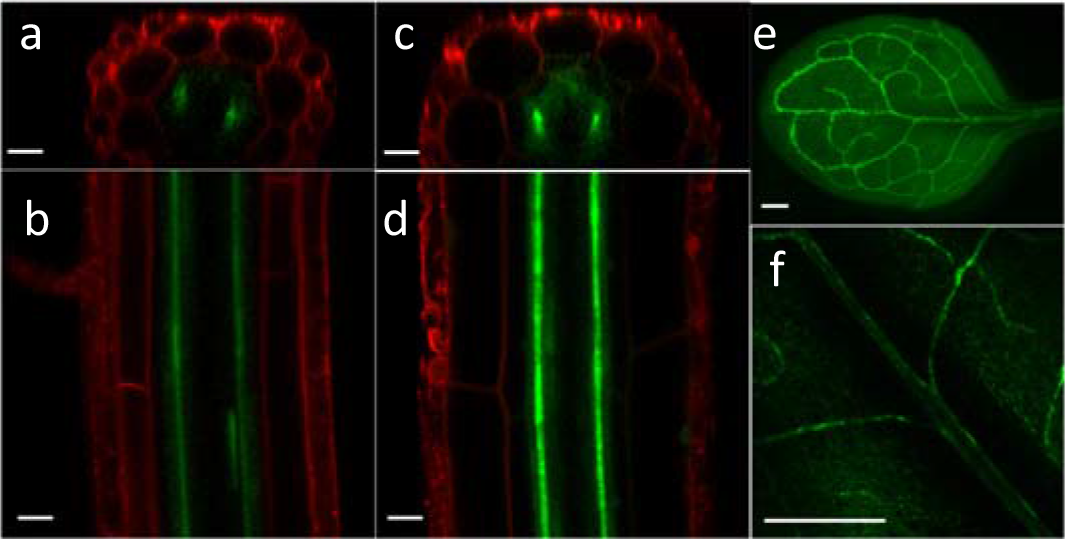
Expression pattern of IMA1 visualized in promIMA1::EYFP expressing plants; (a,b) Yellow fluorescence observed in roots of control and (c,d) Fe-deficient plants; (e,f) Expression of IMA1 in leaves. Scale bar = 30 μm for roots, 500 μm for leaves.

Next, we reciprocally grafted IMA1 Ox shoots onto wild-type rootstocks (IMA1 Ox/WT) and wild-type shoot scions onto IMA1 Ox rootstocks (WT/IMA1 Ox), and determined the FCR activity of the graft combinations. The *opt3-2* mutant was used as a positive control. Under Fe-replete conditions, the wild type displayed low FCR activity, whereas *opt3-2* and IMA1 Ox plants displayed high FCR rates (Fig. 6a). Wild-type scions grafted onto *opt3-2* rootstocks had low FCR activity whereas *opt3-2/*WT grafts displayed high FCR, confirming that altered signaling from the shoot is causative for the constitutive root Fe-deficiency response of the *opt3-2* mutant. WT/IMA1 Ox grafts exhibited high root FCR activity, suggesting that increased IMA levels in roots are sufficient to trigger a constitutive Fe-deficient response. Wild-type rootstocks grafted with IMA1 Ox shoots showed a significantly increased FCR when compared to roots of wild-type plants, although the level was somewhat lower than that of *opt3-2* and IMA1 Ox plants. Ungrafted and self-grafted Fe-deficient *ima8x* mutants were unable to induce root FCR (Fig. 4b; Fig. 6b). Reciprocal grafting of the *ima8x* mutant showed that wild-type scions could partly rescue the impaired FCR induction in *ima8x* rootstocks (Fig. 6b), and wild-type rootstocks grafted with *ima8x* scions induced an FCR activity to a level similar to that of self-grafted controls. Furthermore, overexpression of *IMA1* in leaves led to a significant increase of FCR in roots of Fe-sufficient plants. These results support the supposition that IMAs are phloem mobile peptides that positively regulate iron-deficiency response in roots.

**Fig. 6.**
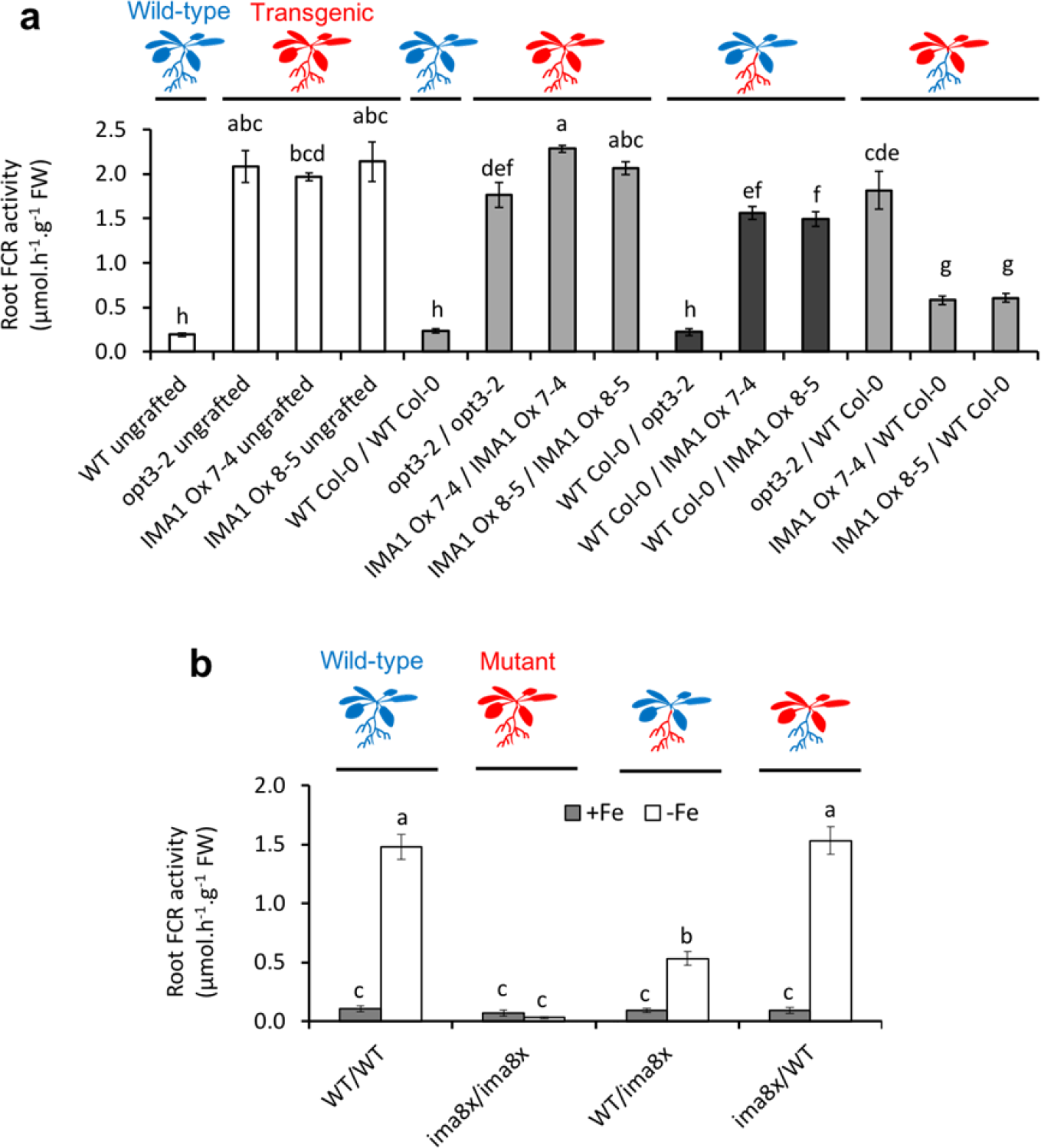
Root FCR activity of reciprocally grafted plants. (a) Grafting of wild-type Col-0 plants, *opt3-2* mutants, and IMA1 Ox lines grown on control (ES) media; (b) Grafting of wild-type Col-0 plants and *ima8x* mutants grown on control and Fe-deficient media. Results are means ± SE (n = 6 sets of 5 roots). Stars indicate significant difference to control plants (Duncan test, *P* ≤ 0.05).

In stable transgenic plants expressing an EYFP:IMA1 fusion protein, fluorescence was observed in the cytosol and nuclei (Fig. 7b,d,f). EYFP:IMA1 lines displayed increased root FCR activity under Fe-replete conditions, indicating functionality of the fusion protein (Fig. 7g). Immunodetection using an anti-GFP antibody revealed a protein of a size between 30 and 40 kDa, consistent with the predicted 34.17 kDa of the EYFP:IMA1 chimera (Fig. 7h). No free EYFP was detected, indicating that the fluorescence signal was representative of the EYFP:IMA1 fusion protein. Interestingly, a similar subcellular localization was observed for the nitrogen signaling protein CEPD1^33^, indicative of putatively similar regulatory mechanisms of the two peptides.

**Fig. 7.**
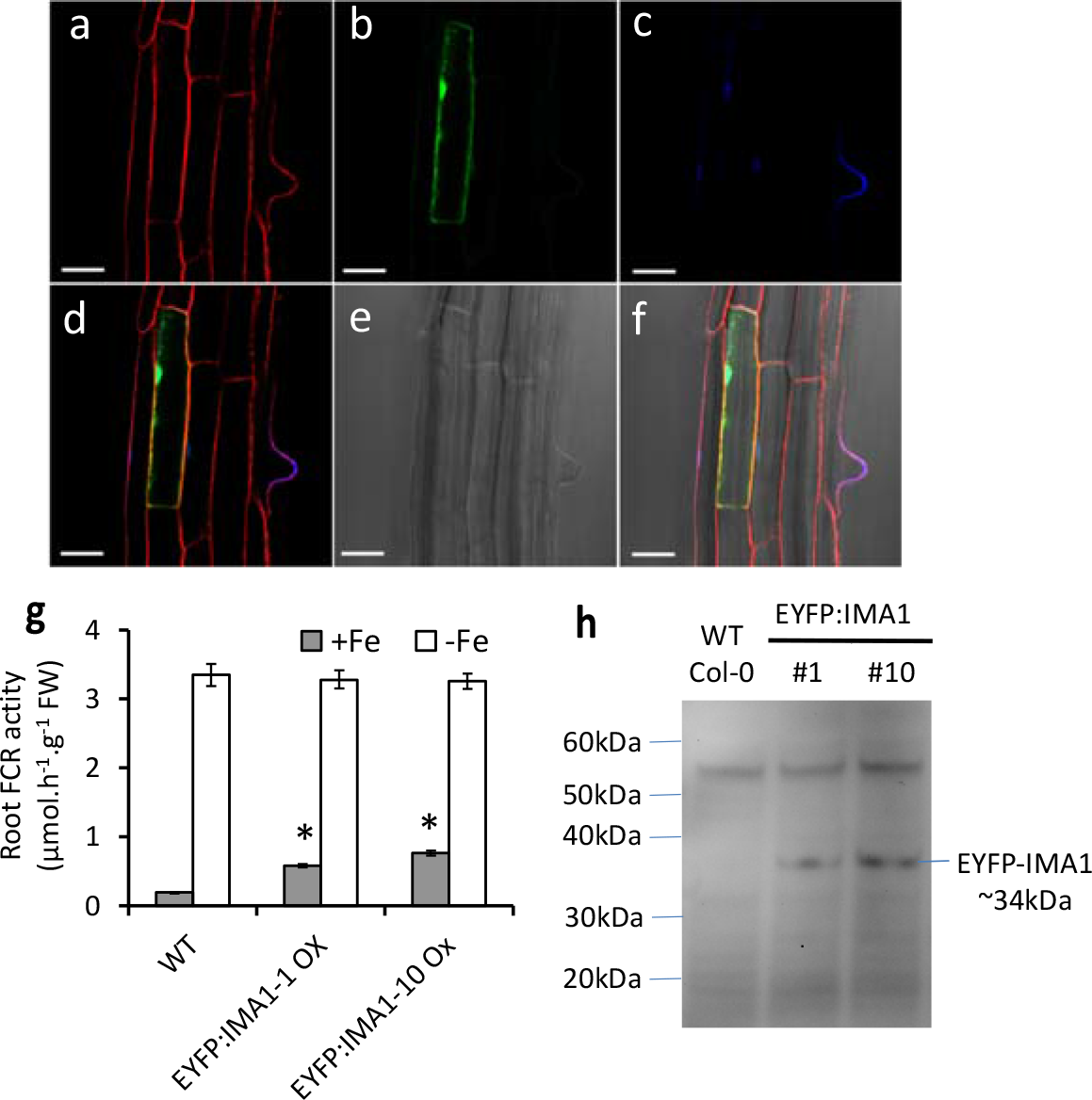
Expression of an EYFP:IMA1 fusion protein. (a-f) Subcellular localization of EYFP:IMA1. (a) Red fluorescence channel showing FM4-64 staining. (b) Yellow fluorescence indicating EYFP localization. (c) Blue fluorescence revealing nuclei stained with DAPI. (d) Merge of the three fluorescence channels. (e) Bright field image. (f) Merge of all channels. (g) Root FCR activity of transgenic plants overexpressing EYFP:IMA1 fusion protein. (h) Western blot with anti-GFP antibodies. Results are means ± SE (n = 6 sets of 5 roots). Stars indicate significant difference to control plants (Duncan test, *P* ≤ 0.05). Scale bar = 30 μm.

Systemic Fe signaling was hypothesized to be mediated by cycling Fe through the phloem, acting as a repressive signal on root Fe uptake^31^. The aspartic acid stretch in the IMA motif is likely to exhibit affinity for metal ions. We thus investigated whether a synthetic peptide corresponding to the 17 C-terminal residues of IMA1 (IMA1pep) could form metal complexes using ESI-MS. Mass spectrometry analysis of IMA1pep metal solutions revealed that IMA1pep can bind Fe^2+^, Cu^2+^, Cu^+^, Zn^2+^, Mn^2+^ but not Fe^3+^, forming complexes of up to four metal ions per peptide (Supplementary Fig. 12; Supplementary Table 3). When different metals were provided simultaneously, IMA1pep-Fe^2+^, IMA1pep-Zn^2+^, and IMA1pep-Cu^+^ complexes were observed in the presence of ascorbate as a reductant (Supplementary Fig. 12a); IMA1pep-Mn^2+^ and −Cu^2+^ complexes were only detected under non-reductive conditions (Supplementary Fig. 12b). Only complexes with Fe^2+^ and Mn^2+^were recovered after chromatography (Supplementary Fig. 13a), suggesting that complexes with ferrous Fe and Mn were more stable than other metal/peptide conglomerates. Interestingly, no signal could be detected when peptides were saturated with metal ions and a precipitate formed quickly upon addition of the metal solution (Supplementary Fig. 13b). Together the data suggest that IMA peptides can bind various metal ions with a moderate specificity for Fe^2+^, and saturation of the binding sites destabilizes the protein in aqueous solution.

To investigate if IMA function is conserved across species, we produced transgenic tomato plants expressing *AtlMAl* driven by the CaMV 35S promoter. Fruits of two independent transgenic tomato lines overexpressing *AtlMAl* cDNA were found to contain significantly more Fe, Mn and Zn than control plants (Fig. 8a). Perls’-DAB staining revealed pronounced Fe accumulation in the transgenic plants that was confined to the vasculature (Fig. 8b). To prove whether heterologous expression of an *IMA* ortholog from the Strategy II plant rice produces the same phenotype than *AtIMAs*, we expressed *OsIMA1* cDNA in *Arabidopsis.* In rosettes of OsIMA1c Ox 12-2 and OsIMA1c Ox 7-1, the Fe concentration was increased by 1.5 to 2-fold, respectively, when compared to wild-type plants (Fig. 8c). Increased Fe levels in rosettes of the transgenic plants were confirmed by Perls’ staining and were absent in control plants (Fig. 8d). Under Fe-sufficient conditions, roots of OsIMA1 Ox lines showed a slight but significant elevation in FCR activity relative to wild-type plants, indicating that the accumulation of Fe was associated with an induced Fe deficiency response in roots (Fig. 8e).

**Fig. 8.**
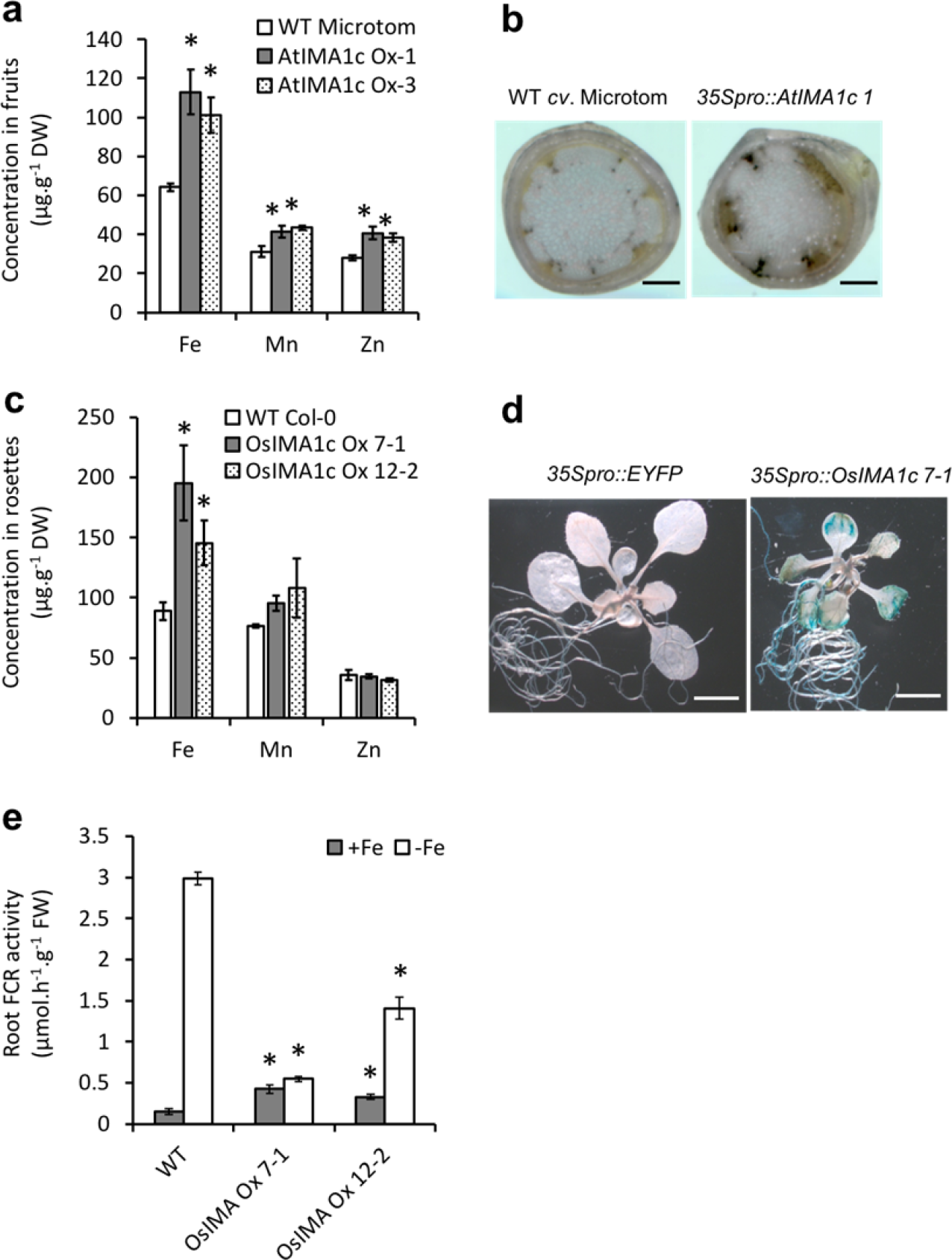
IMA function is conserved across species. (a) Fe, Zn and Mn concentrations in fruits of transgenic tomato plants (n = 3 sets of 3 plants) expressing 35Spro::AtIMA1_cDNA_ (AtIMAlc Ox). (b) Visualisation of Fe by Perls’-DAB staining in cross-sections of stems from wild-type (*cv*. Microtom) and AtIMA1c Ox plants, scale bar = 100 μm. (c) Fe, Zn and Mn concentrations in rosette leaves of *Arabidopsis* plants (n = 3 sets of 15 plants) expressing 35Spro::OsIMA1_cDNA_ (OsIMA1 Ox). (d) Root FCR activity of OsIMA1 Ox plants. (e) Visualisation of Fe by Perls’ staining in seedlings of OsIMA1 Ox plants, scale bar = 300 μm. Results show means ± SE. Stars indicate significant difference to the control plants grown under the same conditions (Duncan test, *P* ≤ 0.05).

## Discussion

We here report on the identification of a novel family of highly Fe-responsive peptides that share a short bipartite C-terminal sequence motif which is critical for Fe uptake. A database search revealed that genes encoding IMA peptides are present in all angiosperms for which data are available, indicating conservation of the motif among flowering plants. Despite their high expression levels under Fe-deficient conditions, IMA peptides have not been recognized as a family due to several constraints that render the identification of a shared consensus sequence difficult. BLAST searches for IMA peptides are hampered by their highly variable N-termini, the presence of an Asp stretch of low complexity which masks the motif for search algorithms, a gap with variable amino acids in the middle of the motif, and the high variability of the ORF size, ranging from 23 to 86 amino acids. Interestingly, partial IMA motifs were also found at the C-terminus of rirA, the main regulator of Fe uptake of plant-interacting alphaproteobacteria such as *Agrobacterium tumefaciens*^47^, as well as in the TonB Fe-siderophore receptor of *Streptomyces* bacteria.

The *Arabidopsis* genome contains eight *IMA* genes, six of which were suggested to produce non-coding RNAs (IMA3-8)^48,49^, a prediction based on the small size of the ORF and the absence of orthologs using BLAST. The high conservation of the IMA consensus amino acid motif and the functionality of the IMA motif when expressed without the non-conserved N-terminal part of the peptide strongly support a function of IMAs at the peptide level. This assumption is corroborated by the observed translation of *IMA1* and *IMA3* mRNA in genome-wide ribosome profiling surveys^46,50^ (Supplementary Fig. 14). Endogenous IMAs have, however, neither been detected through antibodies nor by mass spectrometry. This could be explained by an inherent instability of the peptides and/or low translation of the transcript; stretches of aspartic acid residues were shown to negatively impact translation^51^. Direct evidence for a function of IMAs as peptides derives from the observed increase in FCR activity in plants expressing an IMA1:EYFP fusion protein.

Although the exact molecular mechanism by which IMAs regulate Fe uptake genes remain to be elucidated, binding to Fe^2+^ may control the stability of IMA peptides and thereby regulate Fe uptake. Negatively charged residues such as aspartate participate in the Fe coordination in numerous proteins^52,53,54^. Consistent with the putative metal-binding properties of the Asp stretch in the IMA motif, we showed that IMA peptides can bind Fe^2+^ and other metals. Saturation of the binding sites triggered precipitation of the peptide. Because IMAs are predominantly expressed under low Fe availability, we hypothesized that the instability of IMAs constitute a negative feedback on the Fe uptake machinery which is triggered by phloem Fe under Fe-replete conditions. This would explain the very low accumulation of EYFP:IMA1 protein in the overexpressor, the difficulty to detect endogenous IMA peptides, as well as the relative moderate effect of IMAs from wild-types scions on *ima8x* rootstocks FCR.

The massive induction of IMA-encoding genes in response to Fe deficiency observed in several angiosperms suggests that the function of peptides of the IMA family in Fe homeostasis is conserved across species. The expression of *IMAs* is not regulated by Cu^55^ or Zn deficiency^56^ and is highly correlated with several well-established Fe-specific regulatory genes^17^. Overexpression of the IMA motif overrides the repression of Fe uptake exerted by an adequate Fe status of the plant and triggers an Fe deficiency response in root cells, leading to an increase in the concentration of primary and secondary substrates of the high affinity Fe transporter IRT1 (*i.e.* Fe^2+^, Mn^2+^ and Zn^2+^) in roots and aerial plant parts. On the other hand, silencing of all the *IMA* genes in the *ima8x* mutant leads to lethality in absence of a drastic Fe supplementation and impairs the response to Fe deficiency. It thus appears that IMAs represent an integral component of cellular Fe homeostasis, which is not confined to taxa that have adopted a reduction-based (i.e. Strategy I type) Fe acquisition system such as *Arabidopsis*. Our data further show that the level of IMA peptides dictates the uptake of Fe by acting upstream of the species-specific Fe acquisition machinery. The strong phenotype of *ima8x* mutants show that functional IMAs are crucial for cellular Fe homeostasis under both Fe-replete and Fe-deficient conditions. The excess Fe phenotype of IMA Ox lines is reminiscent of Fe over-accumulating mutants defective in shoot-to-root signaling such as *opt3*^30–31^ and the pea mutant *dgl*^28^, supporting a putative role for IMAs as a promotive signal in the inter-organ regulation of Fe uptake. IMA functionally resembles the role of CEPD1/CEPD2 in systemic nitrogen signaling^33^, suggesting that peptides may be critical in orchestrating the demand of the plant to tune the uptake of mineral nutrients from the soil.

## Material and methods

### Plant growth conditions

Seeds of *Arabidopsis thaliana* (L.) Heynh, ecotype Columbia (Col-0), were surface sterilized and germinated on media containing KNO_3_ (5 mM), MgSO_4_ (2 mM), Ca(NO_3_)_2_ (2 mM), KH_2_PO_4_ (2.5 mM), H_3_BO_3_ (70 μM), MnCfe (14 μM), ZnSO_4_ (1 μM), CuSO_4_ (0.5 μM), CoCl_2_ (0.01 μM), Na_2_MoO_4_ (0.2 μM), and FeEDTA (40 μM), solidified with 0.4% Gelrite pure (Kelco), 1.5% sucrose and 1 g/L MES (ES media^57^). The pH was adjusted to 5.5 with KOH. Seeds were sown on Petri plates and stratified for 2 days in 4 °C in the dark before being transferred to a growth chamber and grown at 21 °C under continuous illumination (50 μmol m^−2^ s^−1^). Standard ES media was supplemented with either 40 μM FeEDTA (+Fe plants), 400 μM FeEDTA (400 Fe plants), or without Fe and 100 μM 3-(2-pyridyl)-5,6-diphenyl-1,2,4-triazine sulfonate (−Fe plants). Non-available Fe (navFe) plants were grown on ES media at pH 7 buffered with 1 g/L MOPS and with 10 μM FeCl_3_. Seeds were germinated and grown for 13 days on the respective media.

Grafting of 5 days-old seedlings was performed using a collar-and hormone-free method described in^58^. For *ima8x* mutants, plants were grown on ES media containing 200 μM Fe-EDTA prior to grafting.

For elemental analysis of seeds and leaves, plants were grown on media for 13 days as mentioned above, transferred to soil containing peat moss (Jiffy), perlite (Rover Green Agriculture Co. Ltd.), and King Root Plant Medium #3 (Rover Green Agriculture Co. Ltd.) at a 10:1:1 ratio, and placed in chambers at 22° C with a photoperiod of 16 hours light and 8 hours darkness at a light intensity of 100 μmol.m^−2^.s^−1^. For seed harvest, Aracons (BETATECH bvba, Ghent, Belgium) were placed over plants a week after bolting. Pots were individually watered twice a week with 50 to 100 mL of tap water and fertilized with ES nutrient solution at the 4 to 6 leaf stage and during bolting.

### Generation of transgenic lines

Full-length *AtIMA1* cDNA was amplified with engineered BamHI sites and cloned into BamHI digested and de-phosphorylated pBIN-pROK2 (Arabidopsis Biological Resource Center) to generate the pROKIMA1 binary vector, which was used for *Arabidopsis* (lines IMA1c Ox 0-8, 1-4, 2-1 and 3-4) and tomato transformation (lines AtIMA1c Ox 1 and 3). For constructs used for the overexpression of *AtIMA1* (lines IMA1o Ox 7-4 and 8-5), *IMA1oΔ1*, *IMA1oΔ2*, *IMA1oΔ3* and *IMA3*, the ORFs were cloned into PCR8/GW/TOPO (ThermoFisher Scientific). ORFs were subsequently transferred into the pH2GW7 vector^59^ (obtained from the Vlams Instituut voor Biologie) by Gateway™ LR recombination, yielding the pHIMA1 and pHIMA3 vectors. *IMA1* deletions were generated by PCR using pIMA1TOPO as a template and recombined with pH2GW7 to produce the binary pHIMA1Δ1, pHIMA1Δ2 and pHIMA1Δ3 vectors. For OsIMA1 Ox, EYFP Ox, and IMA1pep lines, the full-length cDNA of gene *LOC_Os()1g45914*, the EYFP ORF, and the partial IMA1 ORF encoding the last 17 amino acids with an engineered upstream ATG codon, respectively, were subcloned into the pENTR™/D/TOPO vector and recombined with pH2GW7 by Gateway™ LR recombination to obtain the pHOsIMA1, pHEYFP, and pHIMA1pep vectors. The 35Spro::EYFP:IMA1 construct was obtained by cloning *IMA1* ORF into the PCR8/GW/TOPO vector and subsequent Gateway™ LR recombination with the pGWB542 vector^60^. *Agrobacterium tumefaciens* strain GV3101 (pMP90) was used to transform *Arabidopsis* Col-0 plants via the floral dip method^61^; strain LBA4404 was used to transform the tomato cultivar MicroTom as previously described^62^. All transgenic plants were generated by the Transgenic Plant Core Facility of Academia Sinica. Primers used for cloning are listed in Supplementary Table 4.

### Multiplex genome editing

Target sequences were selected within coding sequences of all eight *IMAs* as close as possible to the ATG. The specificity of the sequences was assessed using the Cas-OFFinder tool (http://www.rgenome.net/cas-offinder/)^63^. Sequences and their target cutting sites are given in Supplementary Table 5. The two cassettes for expression of the multiple gRNA scaffolds were used as described in^64^ with either flanking attL4 and attR1, or attR2 and attL3 recombination sequences, respectively, for the first cassette targeting *IMA1*, *2*, *3* and *7*, and the second cassette targeting *IMA4*, *5*, *6* and *8*. Each cassette was synthesized with its flanking gateway sequences and cloned into a pUC57 vector harboring a kanamycin resistance gene by the Genewiz company (South Plainfield, NJ, USA). The *AtUBQ1pro:SpCas9:tAtUBQ1* cassette from the psgR-Cas9 vector described in^65^ was amplified by PCR using the Phusion II HF DNA polymerase and primers harboring attB1 and attB2 flanking sequences, and cloned into pDONR221 (ThermoFisher Scientific) by BP recombination. The two multiple gRNAs and the Cas9 cassettes were cloned into the pH7m34GW vector^59^ (obtained from the Vlams Instituut voor Biologie) through a LR reaction resulting in the pHCas9IMA8x plasmid. The pHCas9IMA8x vector was subsequently transformed into *A. tumefaciens* GV3101, which was used for *Arabidopsis* Col-0 plants transformation using the floral dip method as described above. In total, 190 transformed plants were selected on media containing hygromycin, and fragments of 1 to 1.5 kb surrounding each *IMA* gene were sequenced. Several plants exhibited mild to severe chlorosis. The most severely affected plants harbored either deletions, frameshifts, or sequence modifications, leading to complete disruption of all eight *IMA* genes. Deletions and mutations in the *ima8x* mutant were identified by PCR and confirmed by sequencing. The 35Spro::IMA1_ORF_ and 35Spro::EYFP:IMA1 constructs described previously were transformed into the *ima8x* mutant background.

### Ferric chelate reductase activity

Ferric chelate reductase activity was measured as described in^66^ using roots from 5 to 10 seedlings (10-25 mg fresh weight) at the 4 to 6 leaf stage. Plants were incubated for 1 h in the dark with mild shaking in 2 mL assay solution consisting of 100 μM Fe^3+^EDTA, 300 μM bathophenanthroline disulfonate (BPDS) in 10 mM MES at pH 5.5. Fe^2+^-BPDS_3_ concentration was determined by reading the absorbance at 535 nm on a PowerWave XS2 plate reader (BioTek Instruments, USA). Experiments were conducted at least three times independently.

### Determination of mineral concentrations

Roots and shoots from 3-week-old wild-type and AtIMA1c Ox plants grown under control conditions were harvested separately. Mineral nutrient analysis was determined by inductively coupled plasma mass spectrometry (ICP-MS). Five plants were harvested per treatment and genotype, dried in a conventional oven at 60 °C, and ground in a stainless-steel mill. Aliquots (~0.1 g dry weight) were digested in 65% HNO_3_ and diluted to 14 mL final volume in MilliQ water prior to analysis with a 7700x ICP-MS (Agilent). For analysis of Fe only, plants were dried in an oven at 60 °C, mineralized with 225 μL 65% HNO_3_ at 96 °C for 6 h, and oxidized with 150 μL 30% H_2_O_2_ at 56 °C for 2 h. Fe concentrations were calculated from A_535nm_ of an assay solution that contained 1 mM BPDS, 0.6 M sodium acetate and 0.48 M hydroxylamine hydrochloride against a standard curve made with FeCl_3_.

### Biomass and chlorophyll measurement

For biomass determination, rosettes of about twenty 13-day-old seedlings were weighted immediately after harvest. Subsequently, seedlings were ground with a TissueLyzer bead mill and chlorophyll was extracted in 80% acetone. Total chlorophyll was calculated from absorbance measured at 645, 662, and 750 nm with a PowerWave XS2 plate reader (BioTek Instruments, USA).

### Perls’ staining for Fe(III)

*Arabidopsis* seedlings were vacuum infiltrated with Perls’ solution (2% HCl and 2% potassium ferrocyanide) for 15 minutes and incubated for another 30 minutes. Samples were then rinsed three times with distilled water. For staining embryos and histological sections, Perls’ staining was intensified with diaminobenzidine (DAB) as described in Roschzttardtz *et al*^67^. Subsequently to the staining with Perls’ solution, embryos or slides were incubated for 1 h in a methanol solution containing 0.01 M sodium azide and 0.3% H_2_O_2_, and washed with 100 mM sodium phosphate buffer pH 7.4. Staining was then intensified by 10 min incubation in a solution containing 0.025% DAB, 0.005% H_2_O_2_ and 0.005% CoCl_2_. sections were cut at 5 μm thickness using a RM2255 Leica microtome (Leica, Nussloch, Germany) from rosette leaves of 13-days-old plants embedded in Technovit 7100 resin (Heraeus Kulzer, Wehrheim), and imaged using a Zeiss LSM880 confocal microscope.

### Motif discovery

Sequences of peptides encoded by highly Fe-regulated genes that contain consensus sequence Of motifs were identified in transcriptomes of Fe-deficient *Arabidopsis*^35^, tomato^38^, rice^34^, and soybean^39^ plants. These sequences were used as an input for the MEME suite 4.9.1 online tool^68^. Motif discovery was performed with the Multiple Em for Motif Elicitation tool, and the discovered motifs were then aligned with the input sequences using the Motif Alignment and Search Tool (MAST). The motif was subsequently used for a BLAST^®^ search in the Uniprot database and thorough searches in individual genome databases.

### Gene expression analysis

Total RNA was extracted using the Qiagen RNeasy Plant Mini Kit according to the manufacturer’s instructions. For individual genes, cDNAs were synthesized using the SuperScript III reverse transcriptase (Life Technologies) and real-time qRT-PCR was carried out in an ABI Prism 7500 Sequence Detection System (Applied Biosystems). All qRT-PCR runs were performed as described previously^9^. Primers used for qRT-PCR are listed in Supplementary Table 4. Paired-end stranded RNA sequencing transcriptome analysis of IMA1o Ox 7-4 and 35Spro::EYFP plants was performed as followed. Total RNA of roots and shoots of Fe-sufficient and Fe-deficient 13-day-old plants was extracted with the Qiagen RNeasy Plant Mini Kit. RNA quality was verified using a Bioanalyzer 2100. RIN scores were between 9.8-10 for root and 8.3-8.7 for shoot samples. Libraries were prepared with the TruSeq stranded mRNA LT Sample Prep Kit following the manufacturer’s instructions, and sequenced with HiSeq v4 HT reagents on an Illumina HiSeq-2500 sequencer. Adapter sequences were removed from raw reads using Trimmomatic in keep-two-reads mode before aligning to the *Arabidopsis* TAIR10 genome sequence. For each sample, more than 30 million reads were aligned to TAIR10 gene models with at least 95% identity without indels. Sequences of T-DNAs and non-annotated IMA genes were added to the gene model database. Differential expression analysis was performed with the edge R Bioconductor package using Trimmed Mean of M values (TMM) normalization. For wild-type data, raw data from root^11^ and shoot^35^ were re-analyzed according to the above-mentioned method. Expression levels of genes that were differentially expressed between Fe-deficient and Fe-sufficient wild-type plants were compared to those of IMA1o Ox 7-4 plants grown under similar conditions. RNAseq data of IMA1 Ox and Fe-deficient wild-type Col-0 transcriptomes have been deposited to the Gene Expression Omnibus database and are available under the accession numbers GSE87745 and GSE87760, respectively.

### Synthetic peptide analysis

The IMA1 peptide (ENGGDDDDSGYDYAPAA) was synthesized and HPLC-purified to 98% purity by KareBay™ Biochem Inc. (NJ, U.S.A.). For metal binding assays, peptide solutions were mixed with various metal solutions containing 100 μM Fe, 100 μM ZnSO_4_, 100 μM CuSO_4_, and/or 100 μM MnCl_2_ in 10 mM ammonium acetate buffer at pH 5. Fe was provided either as FeSO_4_ with 500 μM ascorbic acid or as FeCl_3_ without reductant. Complex formation was analyzed for each individual metal and a mix of the four metals using the same method. Peptide binding sites were saturated by addition of 500 μM of each metal to 100 μM of peptide. The mix was immediately injected into a LTQ Orbitrap Elite Hybrid Ion Trap-Orbitrap mass spectrometer (ThermoFisher Scientific) or passed through a TSK Gel amide 80 column.

### Immunodetection and fluorescence imaging

Fluorescence was observed with a confocal Laser Scanning Microscope (Zeiss LSM 510 Meta). Excitation/detection parameters were 514/535-590 nm. Root tissues were ground in liquid nitrogen and proteins were extracted in 2% SDS, 10% glycerol, 60 mM Tris-HCl pH 6.8. Proteins (50 μg) were loaded on a Bis-Tris 4-12% gradient gel (NuPAGE, ThermoFisher Scientific), and blotted onto a PVDF membrane according to manufacturer’s instructions. Immunoblots were performed using a commercial anti-GFP primary polyclonal antibody raised in rabbit (Abcam, ab290) and a secondary anti-rabbit IgG raised in donkey and conjugated to a horseradish peroxidase (GE Healthcare, NA934V).

## Acknowledgments

We thank Thomas J. Buckhout (Humboldt University, Germany) and Marjori Matzke (IPMB, Academia Sinica) for valuable suggestions and critical comments on the manuscript. We further thank Stéphane Mari and Cathy Curie (INRA-SUPAGRO, France) for helpful discussions. We are grateful to Julia Bailey-Serres for kindly providing ribosome profiling data of IMA genes. RNA sequencing was performed by the High Throughput Genomics Core Facility with the assistance of Mei-Yeh Lu, supported by Academia Sinica. We thank Lin-Yun Kuang and Sheng-Ming Chen from the Transgenic Plant Laboratory of IPMB for performing tomato and *Arabidopsis* transformations, Mei-Jane Fang from the IPMB Live Cell Imaging Core Laboratory for the help with confocal imaging, Wen-Dar Lin from the Bioinformatics Core Laboratory at IPMB for bioinformatics support, Yet-Ran Chen and Yu-Chen Huang from the Metabolomics Core Laboratory of the Agricultural Biotechnology Research Center for the support with the ESI-MS. Elemental analysis were conducted through the use of ICP-MS by P.L. supported by the Natural Science Foundation of China (31370280) and the Project of Priority and Key Areas, ISSCAS (ISSASIP1605). This work was supported by an Academia Sinica Investigator Award to W.S.

## Authors contributions

W.S., L.G. and P.L. designed the research, L.G., P.L., W.L. and G.M. performed and analysed experiments, W.S. and L.G. wrote the manuscript.

